# Effects of N361 Glycosylation on Epidermal Growth Factor Receptor Biological Function

**DOI:** 10.1101/2024.07.12.603279

**Authors:** Dennis Lam, Brandon Arroyo, Ariel N. Liberchuk, Andrew L. Wolfe

## Abstract

Epidermal growth factor receptor (EGFR) is a transmembrane tyrosine kinase that is frequently modified by glycosylation post-translationally. In cancer, EGFR amplifications and hotspot mutations such as L858R that promote proliferation have been detected in a significant fraction of non-small cell lung carcinomas and breast adenocarcinomas. Molecular dynamic simulations suggested that glycosylation at asparagine residue 361 (N361) promotes dimerization and ligand binding. We stably expressed glycosylation-deficient mutant EGFR N361A, with or without the oncogenic mutation L858R. Immunofluorescence and flow cytometry demonstrated that the mutants were each well expressed at the cell membrane. N361A decreased proliferation relative to wild-type EGFR as well as decreased sensitivity to ligands. Proximity ligation assays measuring co-localization of EGFR with its binding partner HER2 in cells revealed that N361A mutations increased co-localization. N361A, located near the binding interface for the EGFR inhibitor necitumumab, desensitized cells expressing the oncogenic EGFR L858R to antibody-based inhibition. These findings underline the critical relevance of post-translational modifications on oncogene function.

**SIGNIFICANCE:** EGFR transduces signals from growth factors into cell proliferation and is frequently hyperactivated in tumors. Glycosylation of EGFR at N361 regulates EGFR dimerization, growth factor stimulation of proliferative signaling, and susceptibility to targeted inhibition. Insights into EGFR glycosylation may expand therapeutic opportunities to benefit cancer patients.

## INTRODUCTION

EGFR is a transmembrane tyrosine kinase that transduces extracellular growth factor signals into cellular proliferation. EGFR, also called ErbB1 or HER1, is one of four members of the ErbB family of receptors which includes ErbB2 (HER2), ErbB3 (HER3), and ErbB4 (HER4).^1^ Key domains of EGFR include the extracellular domain, the transmembrane domain, the intracellular kinase domain, and the carboxy tail which is autophosphorylated upon activation (Fig. 1A).^2^ EGFR activity is regulated by binding of agonist ligands, such as epidermal growth factor (EGF) or amphiregulin (AREG), to the extracellular domain, promoting dimerization.^3,4^ The extracellular domain contains a dimerization interface near the ligand binding site.^5^ EGF is a high-affinity binding ligand that is well expressed in multiple organ systems, including lung and breast. AREG is a lower-affinity EGFR ligand, expressed in fibroblast and epithelial systems, which promotes growth less efficiently than EGF.^6^ Ligand binding leads to increased dimerization and autophosphorylation of the intracellular domain of EGFR.^7,8^ This results in stimulation of kinase activity and phosphorylation of direct targets, including EGFR itself in a cross-autophosphorylation event at Y1068, which further activates kinase activity of EGFR to phosphorylate other downstream targets.^9^ This triggers activation of downstream signaling cascades, including Kirsten rat sarcoma virus (KRAS) resulting in phosphorylated ERK, mammalian target of rapamycin complex 1 (mTORC1) resulting in phosphorylated S6, and other growth promoting pathways central to cancer.^1^ These signaling pathways drive cell survival and growth, and cells with hyperactivating alterations in these pathways often become cancerous.

**Figure 1.**
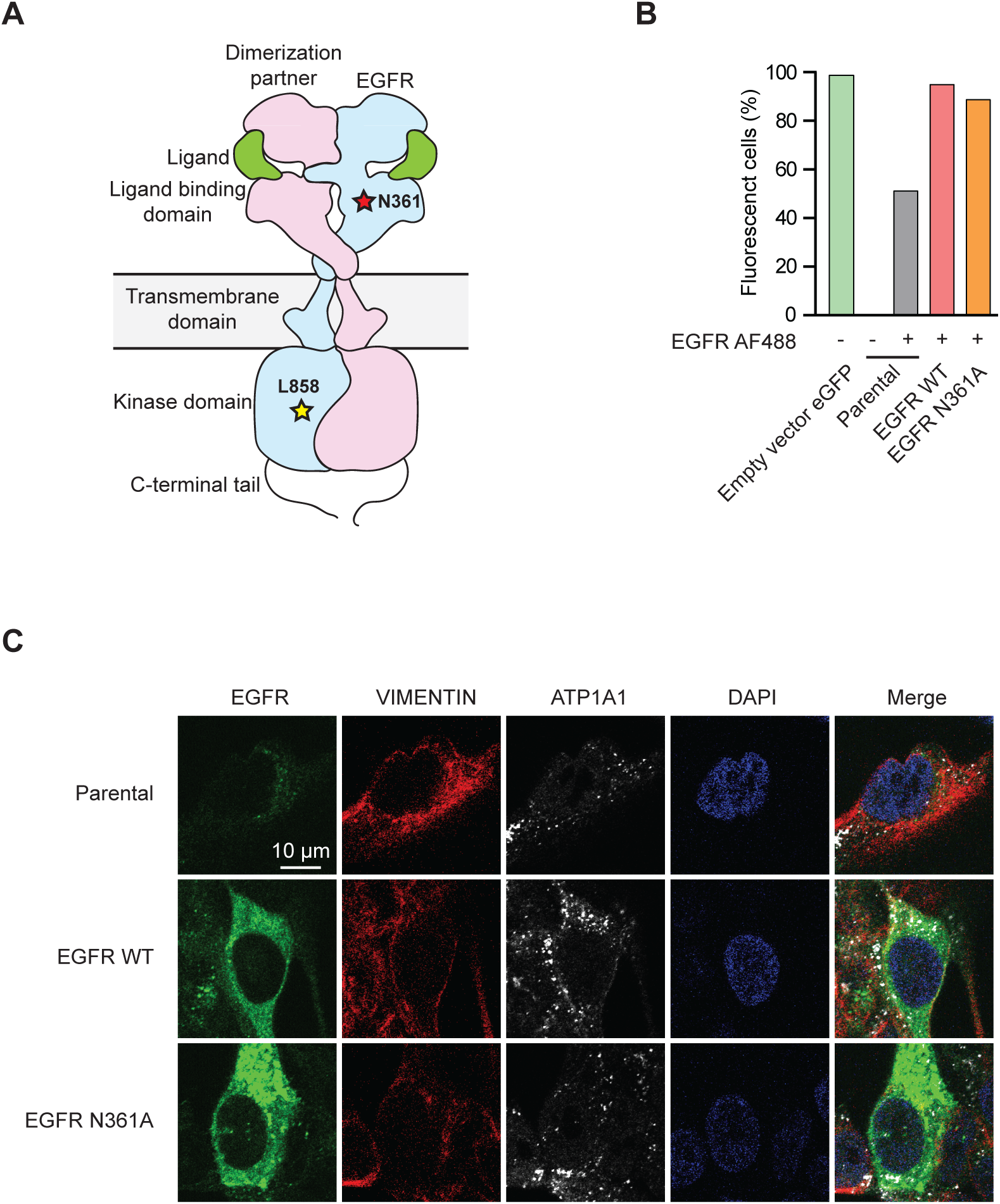
Overexpression of EGFR constructs. **A,** Diagram of domain architecture of EGFR, showing heterodimerization between EGFR and a binding partner such as Her2. Yellow star indicates approximate site of the common oncogenic L858 mutation, red star indicates approximate site of the N361 glycosylation site, green indicates ligands such as EGF or AREG that bind to EGFR to induce dimerization. **B**, Percentage of fluorescent cells from flow cytometry of MCF10A cells that were labeled with anti-EGFR antibody conjugated to Alexa-Fluor 488, or empty puro-IRES-eGFP vector as the stably fluorescent positive control which is not labeled by fluorescent antibody. Cells used were parental MCF10A cells or MCF10A cells overexpressing cDNAs of EGFR wild-type (WT) or EGFR N361A. **C,** Representative immunofluorescent microscopy images of parental MCF10A cells or MCF10A cells overexpressing cDNAs of EGFR wild-type (WT) or EGFR N361A, stained for EGFR, vimentin, ATP1A1, or DAPI.

EGFR is one of the most frequently mutated genes in cancer.^10^ Activating point mutations and amplifications in EGFR are commonly detected in many types of malignancies, occurring in 15-35% of non-small cell lung cancer (NSCLC) and 4-14% of breast adenocarcinomas.^11–13^ In NSCLC, the most common EGFR activating hotspot mutation is a leucine to arginine mutation at residue 858 (L858R) that occurs in 7% of cases.^12,13^ L858 is located in the kinase domain and its mutation to arginine promotes ligand-independent growth, making cells expressing EGFR L858R less dependent on EGF to induce downstream signaling.^14^ Therapeutics inhibiting EGFR kinase activity, such as osimertinib (AZD-92921), are approved for clinical use against EGFR mutant NSCLC. Osimertinib is most effective against EGFR with the kinase domain hotspot mutations T790M and L858R, relative to wild-type EGFR.^15^ Osimertinib is in clinical trials in combination with downstream inhibitors, including antagonists of mutant KRAS.^16^ Several therapeutic antibodies that target the extracellular domain of EGFR have been approved to treat colorectal cancer, NSCLC, and other solid tumors; these include cetuximab, panitumumab, nimotuzumab, and necitumumab. These EGFR targeting antibodies can act by inhibiting dimerization, reducing affinity for ligands, or by attracting inhibitory immune cells.^17–19^

Oncoprotein activity is often regulated by post-translational modifications (PTM). PTMs on receptor tyrosine kinases can modify the ability of ligands to bind, dimers to interact, effectors to be activated, and cells to grow.^20,21^ Glycosylation is a tightly-regulated PTM process where structured sugar groups are attached to target proteins, most commonly on asparagine for *N*-glycosylation and serine or threonine for *O*-glycosylation. Globally, sialylated and fucosylated *N*-glycans are more prevalent in cancer cells, particularly in metastatic cancer cell lines.^22–24^ ErbB family members are heavily glycosylated and several *N*-glycosylation sites have been detected on EGFR.^25–28^ In addition to impacting ligand binding, glycosylation can also can impact the ability of ErbB proteins to be bound by therapeutic antibodies; for instance, disrupting glycosylation of Her2 sensitized the cells to trastuzumab while stabilizing Her2 dimers.^29^ Unglycosylated EGFR N603 mutants have increased propensity to dimerize without ligand, but are unable to increase cell survival in the absence of stimulatory ligand.^30^

Previous glycoproteomic and total proteomic data revealed a dependency for glycosylation of EGFR at residue N361 (sometimes referred to as N337 corresponding to the structure PDB ID: 7SYE where the 24 amino acids were truncated) on the enzyme uridine diphosphate (UDP)-glucose pyrophosphorylase 2 (UGP2).^31,32^ UGP2 is a key metabolic enzyme that regulates the processing of UDP-glucose, which controls both glycogen production and is a key precursor of *N*-glycosylation post-translational modifications on proteins.^33^ UGP2 is upregulated in cells expressing mutant KRAS and is selectively required in KRAS-driven cancer cells, suggesting that UGP2 is an essential part of the mechanism by which cancer cells reprogram their metabolism.^31^ Knockdown of UGP2 in KRAS-mutant cancer cells decreased EGF/EGFR dependent activation of signaling cascades including two direct targets of EGFR kinase activity.^31^ Knockdown of UGP2 caused decreased phosphorylation of EGFR targets, and this behavior could be disrupted using small molecule antagonists of the EGFR kinase domain, indicating that the mechanism by which depletion of UGP2 caused a decrease in these targets was EGFR-dependent.

Glycosylation at EGFR N361 was previously identified in non-cancerous ovarian lines, lung adenocarcinoma, pancreatic ductal adenocarcinoma, and human epidermoid carcinoma cells.^8,28,31,34–37^ In other ErbB family members, glycosylation at the homologous site to EGFR N361 is conserved in ErbB3 and ErbB4, but not in ErbB2.^28^ N361 sits at the edge of the receptor L domain near the junction with the furin-like cysteine rich domain, forming one face of the extracellular EGF binding cavity.^8,25^ At least three UGP2-dependent glycosylation variants containing mannose and *N*-Acetylglucosamine (GlcNAc) were observed at N361.^31,35^ Mutation of the glycosylation site at EGFR N444 (sometimes referred to as N420) has been implicated in a EGF binding deficiency and defective autophosphorylation of the intracellular domain in response to ligand.^38–40^ N444E stimulated a ligand independent change in tyrosine auto-phosphorylation, while N361E required EGF for tyrosine phosphorylation to be induced.^38^ Molecular dynamic simulations of glycosylation at N361 of EGFR showed large changes in side-chains, differences which were magnified in the presence of EGF ligand.^25,26,41,42^ These simulations also suggested that glycosylation sugars at N361 may interact with the opposing EGFR dimer.^25,36^ However, the functional relevance of how glycosylation of EGFR at N361 impacts EGFR protein and cellular behaviors remains unclear.

Here, we stably introduced EGFR mutation incapable of being glycosylated at N361 into non-transformed MCF10A and 293T cell lines. These were selected because they did not have pre-existing mutations constitutively activating the EGFR pathway. We set out to determine how a lack of this specific PTM impacts cell proliferation, subcellular localization of EGFR, ligand binding, inhibitor efficacy, and signaling. Using both in an otherwise wild-type context and in the context of the oncogenic EGFR L858R mutation, we determined that N361A increased co-localization with HER2, yet decreased proliferation and downstream signaling, suggesting non-productive dimerization.

## RESULTS

### Stable overexpression of mutant EGFR N361A

To study the effects of glycosylation at N361 of EGFR, we created the glycosylation-deficient mutant EGFR N361A. We selected non-transformed MCF10A and HEK-293T (“293T”) cells for stable overexpression of EGFR N361A or wild-type EGFR, as these model systems do not have hyperactivation of the EGFR/KRAS pathway by pre-existing cancer-inducing mutations. MCF10A cells normally grow in 20 ng/mL of EGF and express endogenous EGFR, and as a result they are reliant on this signaling pathway. 293T cells express less endogenous EGFR and their media is not typically supplemented with EGF (Supplementary Fig. 1A). We stably transduced MCF10A and 293T cells with wild-type EGFR or EGFR N361A and selected the constructs using puromycin (Supplementary Fig. 1B-C).

The stably transduced MCF10A and 293T cells expressed EGFR cDNAs at similar levels (Fig. 1B-C, Supplementary Fig. 1C-J). Flow cytometry and immunofluorescence of MCF10A and 293T cells showed that stable overexpression of EGFR constructs were significantly higher than endogenous EGFR in non-transduced cells (Fig. 1B and Supplementary Fig. 1B-C,1E-K). Immunofluorescence experiments probing the overexpression and subcellular localization of EGFR demonstrated that our mutant EGFR N361A construct was expressed and was localized to the cytoplasm and cell membrane (Fig. 1C, Supplementary Fig. 2A). In addition, we stably overexpressed in both MCF10A and 293T cells either EGFR L858R or a double mutant cDNA construct with both the EGFR N361A/L858R (Supplementary Fig. 2B-C). Both the L858R and N361A/L858R double mutant were detected by flow cytometry to be well-expressed (Supplementary Fig. 1D). In cells expressing these constructs, both L858R and N361A/L858R retained similar subcellular localization to wild-type and N361A. (Fig. 1C). These results indicated that N361A substitution did not cause significant protein mis-localization in either an the EGFR wild-type or L858R background.

### Differential effects of N361A on proliferation in response to ligand stimulation

To determine the effects of glycosylation at N361 on cell proliferative responses, including responses to natural agonist ligands, we compared the growth of our stably transduced cell lines in media that contained no growth factors (“starved media”), normal media, and media that contained 20 ng/mL more EGF (“stimulated media”). Overexpression of EGFR WT in MCF10A cells in EGF-stimulated conditions led to accelerated cell proliferation relative to empty vector (Fig. 2A). However, overexpression of EGFR N361A did not have the same effect and caused a significantly smaller increase in growth in stimulated conditions (Fig. 2A). This phenotype was not observed in normal or starved media conditions (Supplementary Fig. 3A-B). Growth of all cell genotypes in starved media equally mitigated proliferation (Supplementary Fig. 3B, 3E). Unlike MCF10A cells, 293T cells do not normally depend on the EGF ligand in the media to stimulate the EGFR pathway to promote growth, and thus addition of EGFR constructs did not accelerate 293T proliferation (Supplementary Fig. 3C-E).^43^ In normal media, cells with double mutant EGFR N361A/L858R proliferated less than L858R alone in both MCF10A and 293T cells (Supplementary Fig. 4).

**Figure 2.**
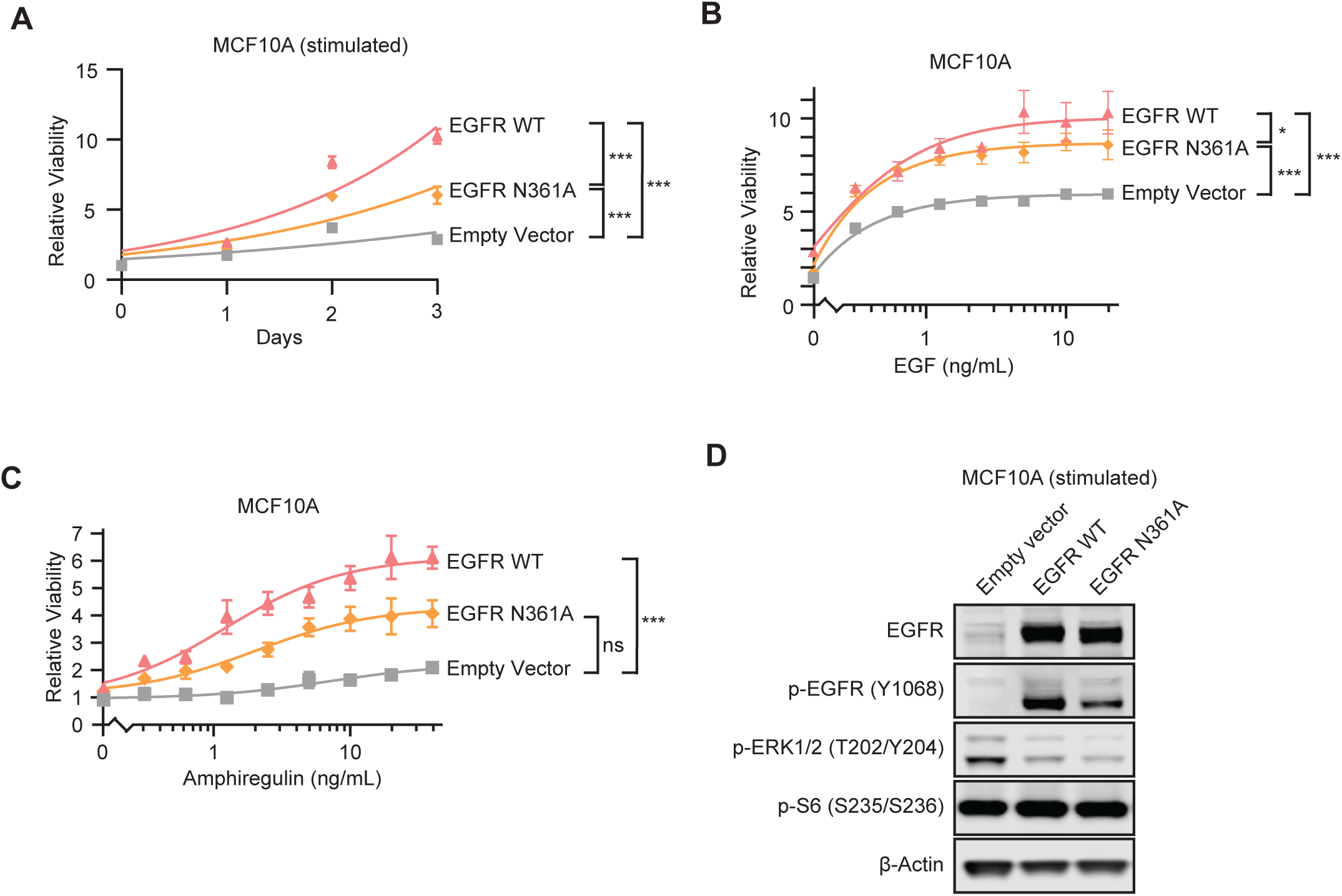
N361A reduces proliferation and response to ligands. **A,** Time course of relative viability of in parental MCF10A cells or MCF10A cells overexpressing cDNAs of EGFR WT or EGFR N361A, measured by Cell-Titer Glo luciferase assay (CTG) in stimulated media with 20 ng/mL of EGF. **B,** Dose course of relative viability of MCF10A cells overexpressing the indicated constructs after stimulation by EGF for 72 hours, measured by CTG. **C,** Dose course of relative viability of parental MCF10A cells or MCF10A cells overexpressing cDNAs of EGFR WT or EGFR N361A upon stimulation by AREG for 72 hours, both measured by CTG; ns = not significant, * p < 0.05; *** p < 0.001, n = 3; **D,** Immunoblots of whole cell lysates of parental MCF10A cells or MCF10A cells overexpressing cDNAs of EGFR WT or EGFR N361A stimulated with an additional 20 ng/mL of EGF for 15 minutes.

We next tested whether the N361A mutation affected the ability of EGFR to respond to growth-stimulating ligand signals with a series of dose course experiments. In 72-hour EGF ligand dose course viability assays, cells with EGFR N361A were deficient at converting higher concentrations of EGF into increased proliferation, relative to cells with EGFR WT (Fig. 2B). Similarly, in a dose course of the alternative EGFR ligand AREG, at high AREG concentrations EGFR WT significantly increased growth relative to empty vector, but EGFR N361A did not (Fig. 2C). In MCF10A cells, L858R and double mutant L858R/N361A were not significantly different at responding to EGF; however, AREG-stimulated MCF10A cells expressing the double mutant EGFR L858R/N361A were able to proliferate significantly faster than EGFR L858R (Supplementary Fig. 6A-B). In 293T cells, which do not normally depend on EGF for sustained viability, there was no dose-dependence response to ligand observed; however, 293T cells overexpressing the double mutant N361A/L858R had slightly less viability in response to AREG relative to L858R (Supplementary Fig. 5A-B, Supplementary Fig. 6C-D). These results show that N361A caused growth deficiencies and decreased proliferative response to ligands.

### N361A reduced cell signaling via the EGFR pathway

To investigate the effects of loss of glycosylation at EGFR N361 on downstream cellular signaling, we performed immunoblots on cell lysates following a brief stimulation with EGF (Fig. 2D and Supplementary Fig. 7). In MCF10A cells, total EGFR overexpressing cDNAs of EGFR WT and N361A transduced cells were both higher than that of empty vector, confirming that our overexpression was successful (Fig. 2D). The cross-autophosphorylation site p-EGFR Y1068, a marker of activation, was more elevated in MCF10A and 293T cells overexpressing EGFR WT than EGFR N361A, (Fig. 2D and Supplementary Fig. 7A). N361A also caused a decrease in downstream p-ERK relative to WT in MCF10A cells. In cells with L858R or N361A/L858R, we again observed comparable overexpression of total EGFR, but the double mutant had elevated p-EGFR and p-ERK relative to L858R (Supplementary Fig. 7B-C). Phosphorylation of S6, a marker for mTORC1 pathway activation, was unchanged in all lines (Fig. 2D, Supplementary Fig. 7). These results are consistent with N361A causing a RAS/ERK pathway signal transduction deficiency.

### Co-localization of EGFR and Her2

Glycosylation can regulate heterodimerization of receptor tyrosine kinases.^44^ To evaluate the impact of the glycosylation-deficient N361A mutant on co-localization of EGFR and one of its binding partners Her2, we performed proximity ligation assays on the MCF10A cell line panel in the presence of EGF, an agonist ligand that promotes dimerization between EGFR and Her2.^45^ Interestingly, EGFR N361A cells had strongly elevated PLA intensity per cell relative to EGFR wild-type or negative unstained controls, indicating that N361A drives increased co-localization (Fig. 3). Concordantly, the EGFR N361A/L858R double mutant displayed significantly more co-localization than L858R alone (Supplementary Fig. 8A-B). While co-localization in the glycosylation deficient mutant is increased, it is unclear from the PLA assay alone whether the co-localization represents productive dimerization, or if the N361A mutation acts as a dominant negative with respect to the function of the heterodimeric complex.

**Figure 3.**
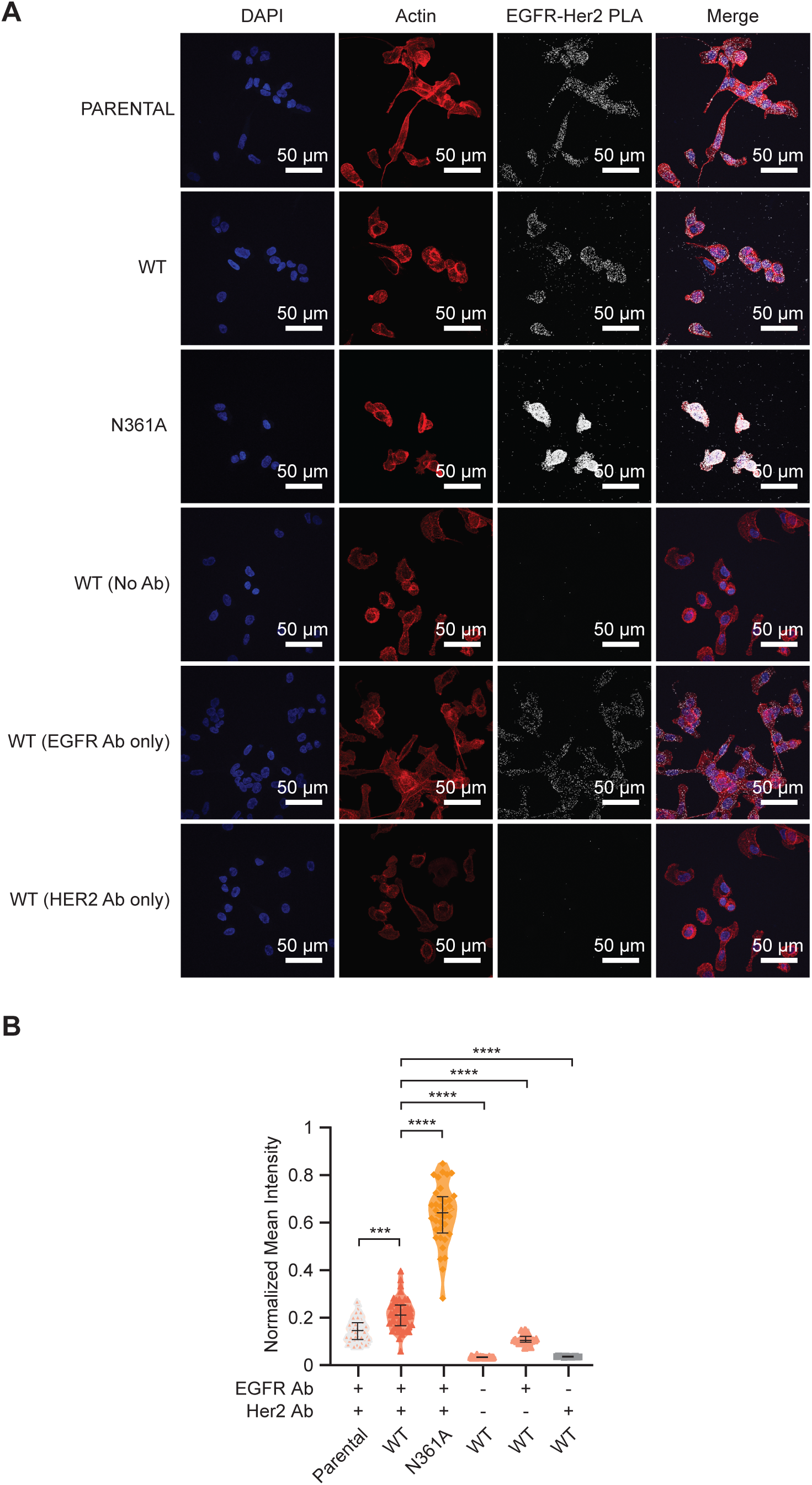
**A,** Immunofluorescent images of MCF10A cells from *in situ* proximity ligation assays using primary antibodies against EGFR and HER2 to assess co-localization. Cells are parental or overexpress exogenous cDNAs of EGFR WT, EGFR N361A, EGFR L858R, or a single construct containing two mutations EGFR N361A/L858R. Scale bar represents 50 μm. **B,** Representative quantification of mean pixel intensity of cellular regions defined by rhodamine Actin stain, including five replicate images per condition. * p < 0.05; ** p < 0.01; *** p < 0.001; **** p < 0.0001, n = 5.

### N361A desensitizes cells to inhibition of EGFR with extracellular antibody necitumumab

To assess the impact of N361A on antibody inhibitors of the extracellular domain of EGFR, we used necitumumab, a clinically approved antibody inhibitor that targets domain III of EGFR.^18^

Ten-point necitumumab dose course treatments from 0 to 30 μM demonstrated that while there was significant growth inhibition by the antibody inhibitor necitumumab of MCF10A cells expressing wild-type EGFR (IC50 of 2.6 μM), the cell line expressing N361A was completely insensitive to the antibody inhibitor, much like the vector control (Fig. 4A). There was no significant inhibition between 293T cells expressing either wild-type EGFR, EGFR N361A, or vector control (Supplementary Fig. 9A). Accordingly, cells overexpressing EGFR L858R were highly sensitive to necitumumab (MCF10A IC50 4.0 μM, 293T IC50 4.5 μM), but in both MCF10A and 293T cells, the lines overexpressing the double N361A/L858R mutant were completely insensitive at all tested doses of necitumumab (Supplementary Fig. 9B-C). This striking phenotype suggests that the glycosylation site N361 is required to be intact for necitumumab efficacy.

**Figure 4.**
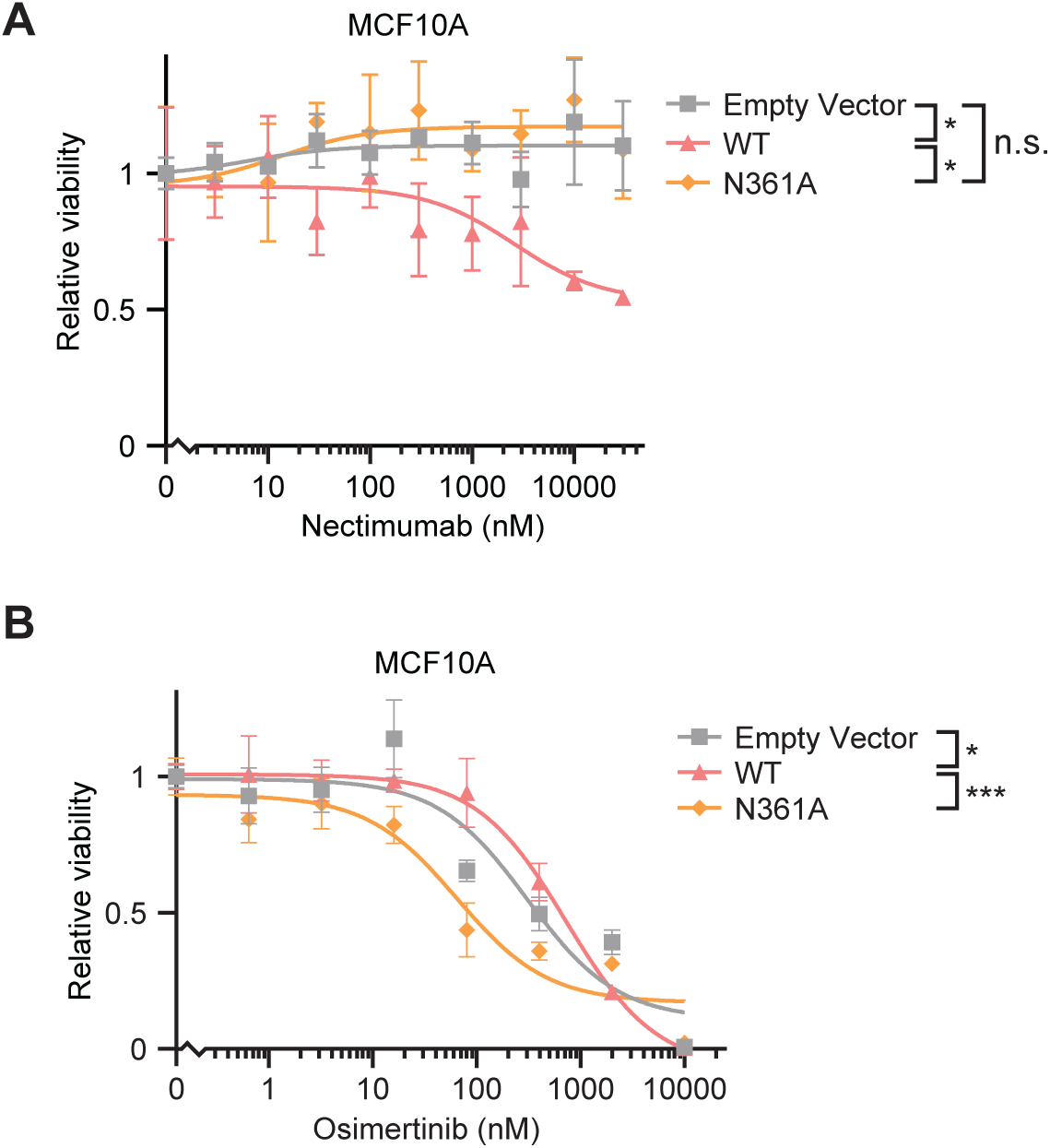
**A-B,** Dose course of relative viability measured by CTG of parental MCF10A cells or MCF10A cells overexpressing cDNAs of EGFR WT or EGFR N361A after 72 hours of treatment with either **(A)** necitumumab or **(B)** osimertinib. ns = not significant, * p < 0.05; *** p < 0.001, n = 3.

Knowing that drug response of EGFR may be dependent on its glycosylation state, we evaluated the response of cells expressing EGFR N361A to the tyrosine kinase domain inhibitor osimertinib, which is used clinically against NSCLC with EGFR L858R mutations or exon 19 deletions.^15^ N361A sensitized MCF10A cells to inhibition by osimertinib than wild-type EGFR (WT IC50 728.5 nM, N361A IC50 64.8 nM) (Fig. 4B). This difference was not observed in 293T cells, which were insensitive to osimertinib at all but the highest of doses (Supplementary Fig. 9E-F).

Consistent with previous reports, we find that MCF10A cells with EGFR L858R mutation are more sensitive to osimertinib than cells with wild-type EGFR (WT IC50 728.5 nM, L858R IC50 361.4 nM) (Fig. 4B and Supplementary Fig. 9D).^46^ MCF10A cells stably expressing the double mutant L858R/N361A were significantly more sensitive to osimertinib than the single mutant L858R cells (L858R/N361A IC50 22.8 nM, L858R IC50 361.4 nM) (Supplementary Fig. 9D). These results further emphasize the importance of considering EGFR glycosylation state when deciding on a treatment course for cancer patients.

## DISCUSSION

Collectively, our data on proliferation, co-localization, and signaling suggest a model where N361A induces formation of dominant non-functional co-localization events.^47,48^ *In situ* PLA experiments showed that the intensity of co-localization events between overexpressed EGFR N361A and endogenous HER2 was approximately threefold higher compared to overexpressed EGFR WT (Fig. 3). Dimerization and activation of EGFR can occur as separate events with different thresholds, and inactive dimers which do not induce kinase activity can form.^47,48^ ErbB receptors may be tuned to be more oncogenic by the upregulation of their *N*-glycan groups.^49,50^ Previous recombinant studies using *in vitro* purified soluble mutant EGFR N361E demonstrated that this mutant was able to dimerize in response to EGF.^38^ HER2 heterodimerization with EGFR induces structural and phosphorylation changes in the phosphorylation of the carboxy tail, which can be impacted by non-functional dimerization.^47,48,51^

Normal heterodimerization between Her2 and EGFR induces phosphorylation of EGFR at Y1068, but cells with a mutation at N361A were deficient for EGFR p-Y1068 (Fig. 2D). Phosphorylation of ERK comes on rapidly within minutes of stimulation of EGFR by ligands like EGF or AREG. EGFR hotspot mutations L858R and T790M increase autophosphorylation, but this activation can be suppressed by EGFR glycosylation (Supplementary Fig. 4B,E).^52^ In the EGFR oncogenic mutant context, glycosylation can also suppress downstream signaling, consistent with our findings in the double mutant cells (Supplementary Fig. 7B-C).^52^ Over time, ERK phosphorylation decreases back to a steady state to prepare cells for future ligand encounters. In our stable cell MCF10A lines, those with constitutive expression of any EGFR construct reached a lower steady state of ERK activation, consistent with adaptation to increased receptor expression. MCF10A and 293T cells overexpressing EGFR N361A were less able to convert new ligand signals into increased p-ERK compared to cells expressing wild-type EGFR, supporting the model that the increased dimerization is non-productive (Fig. 2D, Supplementary Fig. 7A). The stable mutant did not measurably impact p-S6, indicating that disruption of glycosylation at N361 primarily impacts the RAS axis, while sparing mTOR signaling. In summary, *N*-glycosylation at EGFR N361 is important for integrating ligand signals into cellular proliferation.

Structurally, N361 is located in the extracellular domain proximal to the EGF binding site, which is formed from the cavity of Domains I through III.^8^ Both EGF-ligand binding and *N*-glycosylation of EGFR stabilize and place Domain III monomers closer together.^44^ In molecular dynamic simulations, glycosylation groups on EGFR N361 are predicted to bind to EGFR residues 329–333, which are directly involved in electrostatic binding to dimerization partners.^25^ Glycosylation at N361 may impact the structure of this domain leading to an increased affinity for EGF. Consistent with a model where N361A mutation negatively impacts EGF binding affinity, the alanine mutant at N361 resulted in decreased efficacy of EGF on promoting growth of MCF10A cells relative to wild-type, and the double mutant of N361A and L858R was less sensitive to EGF than the single L858R mutant at low concentrations. This would be consistent with a model where glycosylation at N361 shifts the receptor L domain into a conformation with a higher affinity for ligand binding.

Other glycosylation sites such as N151, N328, and N603 on EGFR, and the choice of ligand, may impact the how ligand binding domain acts to stabilize EGFR-EGF binding.^25,36^ Relative to wild-type EGFR, unglycosylated EGFR N603 mutants demonstrated increased propensity to dimerize without ligand, but were deficient at increasing cell survival in the absence of stimulatory ligand.^30^ EGF has higher affinity binding for EGFR than AREG, and as expected stimulation with EGF led to larger increases in proliferation than AREG.^53,54^ (Fig. 2B-C). EGF is more efficient than AREG at stimulating homodimerization of EGFR/EGFR and heterodimerization of HER2/EGFR.^55^ EGF preferentially binds HER2/EGFR heterodimers compared to EGFR homodimers, while AREG has no preference, leading to the selection of EGF-containing media for our PLA experiments (Fig. 3).^55^

This study employed models relevant to normal cell biology and cancer. MCF10A cells were selected because they are grown in EGF-enriched media and have an inherent dependency on EGFR signaling.^43^ In contrast, 293T cells have low expression of EGFR and are not naturally dependent on the EGFR pathway. Our results are consistent with this functional difference, in that MCF10A cells were highly sensitive to osimertinib in an EGFR dependent manner, while 293T cells were only impacted at the highest doses (Fig. 4B and Supplementary Fig. 9D-F). We employed overexpression techniques that do not impact endogenous EGFR, so in cells homozygous for N361A the phenotypes may be even more severe.

Glycosylation of EGFR influences responses to anti-EGFR therapeutic agents. Molecular dynamics analyses of co-crystal structures of EGFR with inhibitory antibody mAb806 found that glycosylation can, in some instances, increase the exposure of the bound epitope.^25^ Mutating away the glycosylation site by introducing N361A may change the EGFR ectodomain structure so that single-agent necitumumab treatment is not able to bind effectively (Fig. 4A). In contrast, kinase domain inhibitors like osimertinib are sensitized by removing glycosylation from EGFR (Fig. 4B).^56^ In ovarian, colorectal, and pancreatic cancer cell lines, overexpression of a glycosylation-promoting enzyme, ST6Gal-I sialyltransferase, increased EGFR activation and enhanced resistance to cell death caused by treatment with an inhibitor of EGFR kinase activity, gefitinib.^56^

Inhibitors of glycosylation are promising tools and may hold a promising future in the clinic. Glycosylation is frequently elevated in tumor cells and is often proposed as a biomarker for cancer detection.^57^ Tunicamycin inhibits UDP-GlcNAc-1-phosphotransferase, an enzyme critical for *N*-glycosylation.^58^ Tunicamycin can decrease EGFR glycosylation, leading to decreased stabilization and reduced activating EGFR phosphorylation.^59^ Tunicamycin decreases signaling through the EGFR/ERK axis more in cancer cells than in normal cells.^59^ Glycosylation subtypes like sialic acid-binding immunoglobulin-like lectin 7 (Siglec-7) can promote immune evasion, leading to the development of Siglec inhibitors.^60^ In screens of lung cancer lines using glycosylation inhibitor NGI-1, lines with genomic alterations in EGFR tended to be among the most highly sensitive.^61^ Use of NGI-1 can overcome acquired resistance to EGFR inhibitors, or can be used effectively as a two-drug combination.^61^ Targeting upstream enzymes that regulate glycosylation, such as UGP2 or Golgi phosphoprotein 3 (GOLPH3), is another promising alternative strategy for cancer.^31,32,62^ EGFR inhibitors have been combined with compounds that can disrupt pre-existing glycosylation or alter global glycosylation patterns – for instance, using erlotinib in combination with 1,3,4-O-Bu3ManNAc to reprogram sialylation in pancreatic cancer.^63^ This study’s finding that the glycosylation-deficient mutant N361A improved the efficacy of EGFR tyrosine kinase inhibitor osimertinib suggests that strategies decreasing glycosylation of EGFR may have clinical relevance at improving the efficacy of EGFR tyrosine kinase inhibitors.

Several unresolved questions about the role of glycosylation at N361 remain. In future studies, it would be illuminating to pursue experiments investigating N361A function in CRISPR knockout lines lacking endogenous EGFR, the roles of N361 in endocytosis and EGFR homodimerization, the effects of N361A on additional phosphorylation sites on HER2 and EGFR, and using CRISPR to create endogenous locus N361A mutant cancer cell lines with or without endogenous activating oncogenic EGFR mutations. ^51,64,65^ Thus, there are many opportunities to employ our knowledge of protein glycosylation to understand cell biology and improve cancer therapy.

## METHODS

### Constructs, cell lines, and time course treatments

Introduction of N361A by site-directed mutagenesis on the pBABE puro EGFR wild-type (EGFR WT, RRID:Addgene_11011) and pBABE puro EGFR L858R (RRID:Addgene_11012) plasmids was performed by Azenta. pBABE puro IRES-eGFP (RRID:Addgene_14430).^66^ All plasmid sequences were independently confirmed by Sanger sequencing.

MCF10A cells were cultured and maintained in DMEM/F12 media (Gibco Cat# 11330057), 5% Horse Serum (Invitrogen Cat# 16050-122) and 1% Penicillin Streptomycin Glutamine (PSG) (Gibco Cat# 10378016), with the addition of 20 ng/ml EGF (Peprotech Cat# AF-100-15), 0.5 mg/mL hydrocortisone (Sigma-Aldrich Cat# H-0888), 100 ng/ml cholera toxin (Sigma-Aldrich Cat# C-8052), 10 μg/ml insulin (Sigma-Aldrich Cat# I-1882), as additional supplements to their complete media. HEK-293T (293T) cells were maintained in DMEM high glucose media (Gibco Cat# 11965092), 10% fetal bovine serum (FBS) (R&D Systems Cat# S11550H) and 1% PSG as their complete media. Cells were maintained at 5% CO2 in a 37°C humidified incubator. Starved media for MCF10A stable cell lines contained maintenance media without horse serum, EGF, or insulin. Starved media for 293T stable cell lines contained complete media without FBS. Stimulated media for both cell lines contained complete media with an additional 20 ng/ml EGF.

All cell lines tested negative for mycoplasma using MycoAlert® Mycoplasma Detection Kit (Lonza Cat# LT07-318). Cell line identities were confirmed using short tandem repeat analysis (ATCC Cat# 135-XV).

For time course experiments, live cells were counted using a Countess 3 Automated Cell Counter and plated at 5,000 cells/well in sterile optical-bottom 96-well plates (Thermo Fisher Scientific Cat# 165306) in 200 μL of starved, normal, or stimulated media for MCF10A and 293T stable cell lines as indicated above. CellTiter-Glo assays were performed at 0 hours, 24 hours, 48 hours and 72 hours after treatment using a SpectraMax i3 (Molecular Devices i3) using CellTiter-Glo Assay kits (Promega Cat# G7572).

### Statistical Analyses

Power analysis was performed to determine the number of replicates needed for the significance level of p < 0.05. Statistical tests are described in figure legends, and all other tests were two-tailed T-tests performed in GraphPad Prism 10 (RRID:SCR_002798) or Microsoft Excel (RRID:SCR_016137). For time- and dose-course experiments, data processing was performed using SkanIt software. Data were normalized to control mock treated conditions at day 0. Two-way ANOVA followed by Tukey’s multiple comparisons test by cell line was performed for EGF growth time course assays. Exponential growth equation was determined using nonlinear fit analysis. Two-way ANOVA followed by Tukey’s multiple comparisons test was performed for EGF and AREG growth dose course assays by cell line using the dose with the greatest divergence. Two-way ANOVA followed by Tukey’s multiple comparisons test was performed for osimertinib and necitumumab growth dose course assays by cell line. IC50s were determined using nonlinear fit analysis, comparing concentration of each inhibitor against response.

Flow cytometry analysis was performed using Flowing Software 2.5.1 (Turku Bioscience) and BD CellQuest Pro (BD Biosciences). One-way analysis of variance (ANOVA) followed by Tukey’s multiple comparisons test was performed to analyze direct-stain flow cytometry. Kruskal-Wallis test followed by Dunn’s multiple comparisons test was performed for EGFR-HER2 *in situ* proximity ligation assay coupled with immunofluorescence. Normalized mean intensity was calculated using CellProfiler Image Analysis Software (RRID:SCR_007358) employing the MeasureObjectIntensity module, which scales pixel intensity within each cell in the image from 0 to 1. For all figures, p-values are shown as: ns represents not significant, * *p* ≤ 0.05, ** *p* ≤ 0.01, *** *p* ≤ 0.001, and **** *p* ≤ 0.0001. Data were plotted in box-and-violin arrangement in GraphPad Prism 10.

See **Supplementary Data** for additional methods information on generation of stable cell lines, dose-course treatments, immunoblot assays, immunofluorescent assays, flow cytometry, and *in situ* proximity ligation assays (PLA) coupled with Immunofluorescence assays.

## Acknowledgements

We gratefully acknowledge Viola Ellison and Leonard J. Ash for advice on immunofluorescence, Nikita Meghani for advice on puromycin selection, Gu Xiao for providing MCF10A cells; Brian Zeglis for use of the LI-COR Odyssey CLx imaging system, Jill Bargonetti for use of the ChemiDoc MP Imaging System and for advice on the manuscript, Leonard J. Ash and Ottavia Bourdain for providing feedback on the manuscript, Shlomo Pallas for modeling of EGFR,Azenta Inc for assistance with mutagenesis and Sanger sequencing; Lloyd Williams from the Bio-Imaging Facility of Hunter College for use of the Nikon A1 Confocal Microscope at Belfer Research Building; Joon Kim from the Flow Cytometry/Genomic Lab of Hunter College for use of the Becton-Dickinson BD FACSCalibur, and thank all members of the Wolfe lab for insights and advice. We acknowledge the Matthew Meyerson lab, the L. Miguel Martins lab, and Didier Trono lab for depositing plasmids in Addgene. This work was supported by Hunter College of the City University of New York (DL, ANL, BA, ALW). Research reported in this publication was supported by the National Cancer Institute of the National Institutes of Health under award number R00CA226363 (DL, ALW). ALW was supported by the National Science Foundation Research and Mentoring Postbaccalaureates (Award 2318923); Temple University Fox Chase Center/Hunter College (TUFCC/HC) Regional Comprehensive Cancer Health Disparity Partnership (U54CA221704); PSC-CUNY Research Award (ENHC55171); and Oncogenuity, Inc. BA was supported by the Maximizing Access to Research Careers Program (MARC, Grant # T34GM149429).

## Authors’ Contributions

**D. Lam:** Conceptualization, data curation, formal analysis, investigation, methodology, supervision, validation, visualization, writing - original draft, writing - review & editing.

**B. Arroyo:** Investigation, validation, writing - original draft.

**A.N. Liberchuk:** Investigation, validation, writing - original draft.

**A.L. Wolfe: Conceptualization,** data curation, formal analysis, funding acquisition, investigation, methodology, project administration, resources, supervision, validation, visualization, writing - original draft, writing - review & editing.

## Data availability

The data and reagents from the study are available in the article and the supplementary files. Plasmids generated in the study are available from Addgene.

