## Supplementary Data and Supplementary Figures for "Effects of N361 Glycosylation on Epidermal Growth Factor Receptor Biological Function"

#### SUPPLEMENTARY METHODS

##### Stable cell lines

Stable MCF10A (RRID:CVCL_0598) and 293T (RRID:CVCL_0063) cell lines overexpressing EGFR constructs or empty vector controls were created. Viral packaging Phoenix-AMPHO (ATCC Cat# CRL-3213, RRID:CVCL_H716) cells were transfected using a 4:1 ratio of plasmid of interest to psPAX2 (RRID:Addgene_12260) in Opti-MEM™ I, Reduced Serum Medium, GlutaMAX™ supplement (Gibco Cat# 51985034) using the Lipofectamine 3000 Kit (Thermo Fisher Scientific Cat# L3000015) following the manufacturer's instructions. After 48-72 hours, supernatants were harvested and 1:1000 polybrene (Santa Cruz Biotechnology Cat# SC134220) was added prior to exposure to MCF10A or 293T cells. An Eppendorf 5810 tabletop centrifuge was used for cell maintenance and assays.

After at least 72 hours of recovery, cell lines were twice selected with 4 µg/ml puromycin (Gibco Cat# A1113803), with each selection lasting one week until untransduced parental control cells were eliminated. To enable antibody-based immunofluorescence, EGFR constructs were chosen that did not contain a fluorescent reporter. An empty vector eGFP control cell line was created and puromycin selected in parallel to the EGFR constructs, and puromycin selection resulted in a greatly increased proportion of GFP expressing control cells by flow cytometry (Supplementary Fig. 1B-C). These stable MCF10A cell lines and 293T cell lines were maintained and cultured in their respective complete media.

Sex as a biological variable did not apply to this study as no human or animal subjects were employed. Both MCF10A and HEK-293T are samples from females.

##### Dose-course treatments

For each cell line, live cells were counted using a Countess 3 Automated cell counter and plated at 5,000 cells/well in sterile optical-bottom 96-well plates (Thermo Fisher Scientific Cat#

165306) in 100 µL of media indicated above. The next day, cells were treated with an additional 100 µL of the indicated final concentration of animal-free recombinant human EGF (Peprotech Cat# AF-100-15), recombinant human AREG (Peprotech Cat# 100-55B), osimertinib (Selleck Chemical LLC Cat# S7297), or necitumumab (Selleck Chemical LLC Cat# A2048). CellTiter-Glo assays were performed at 0 hours and 72 hours after treatment using a Varioskan Lux (Thermo Fisher Scientfic Cat# VLBL00GD2) using a CellTiter-Glo Assay (Promega Cat# G7572). In experiments using osimertinib and necitumumab, all cells received the same final volume of the drug vehicle DMSO.

**Immunoblot assays**

After a 15 min incubation in either normal or stimulated media as indicated above, cells were harvested directly from the tissue culture dish. Cells were removed from plates by scraping, transferred to 15 mL tubes, centrifuged for 300 x g for 5 min, and washed 2 times with 1X ice-cold PBS, then resuspended into 3-fold volume of ice-cold RIPA (Thermo Fisher Scientific Cat# 89901) and 1X HALT (Thermo Fisher Scientific Cat# 78445) buffer in a 1.5 mL centrifuge tube, followed by incubation on ice for 30 min with gentle vortexing every 5 min. These whole-cell lysates were then centrifuged at 14,000 x g for 10 min at 4°C. An Eppendorf 5427R tabletop centrifuge was used for all lysates. Protein concentrations of these cell extracts were determined with a Pierce BCA assay kit (Thermo Fisher Scientific Cat# 23225). Equal amounts of extract per sample were combined with 4X LDS sample buffer to 1X final concentration (Thermo Fisher Scientific Cat# NP0008) and 50 mM dithiothreitol (Thermo Fisher Scientific Cat# R0861), boiled for 70°C for 10 min, and run on NuPAGE™ 4-12% Bis-Tris 1.5mm Mini Protein Gels (Invitrogen Cat# NP0322BOX).

Blot transfers were performed with either a dry-blot or wet-blot transfer system. Dry-blot transfers to iBlot2 Transfer mini nitrocellulose stacks (Invitrogen Cat# IB23002) were performed using an iBlot2 machine (Thermo Fisher Scientific Cat# IB21001) at 25V for 8 min. For wet-blot transfers, protein bands were transferred to 0.2 μm nitrocellulose membrane blots in transfer buffer (2.5 mM Tris, 19 mM glycine, 10% methanol, and 0.1% SDS) at either a constant current of 15 V overnight at 4°C or a constant current of 30 V at 4°C for 1 hour.

Membranes were probed as indicated with the following primary antibodies in 3% bovine serum albumin (BSA) (Sigma-Aldrich Cat# A7906) in 1X TBS-0.1% Tween-20 and its corresponding dilutions in 4°C overnight: 1:5000 β-Actin (Sigma-Aldrich Cat# A5441, RRID:AB_476744), 1:500 EGFR (Cell Signaling Technology Cat# 2232, RRID:AB_331707), 1:500 phospho-EGFR Y1068 (Cell Signaling Technology Cat# 2234, RRID:AB_331701), 1:500 phospho-Erk 1/2 T202/Y204 (Cell Signaling Technology Cat# 9101, RRID:AB_331646), 1:500 phospho-ribosomal protein S6 (p-S6) S235/236 (Cell Signaling Technology Cat# 2211, RRID:AB_331679). Membranes were subsequently washed with 1X TBS-0.1% Tween-20 and probed with these secondary antibodies and its corresponding dilutions at room temperature: 1:10000 anti-mouse 800CW (LI-COR Cat# NC9401842), 1:10000 anti-mouse 680RD (LI-COR Cat# NC0252290), 1:1000 anti-rabbit 800CW (LI-COR Cat# NC9401841), 1:10000 anti-rabbit 680RD (LI-COR Cat# NC0252291). Image processing was performed by LI-COR Odyssey CLx (LI-COR Cat# 9140-09) or ChemiDoc MP (BioRad Cat# 12003154) machine and Image Studio and Image Lab software from LI-COR and Bio-Rad, respectively.

**Immunofluorescence Assay**

For each cell line, live cells were counted using a Countess 3 Automated cell counter and plated at 200,000 cells/well in either clear-bottom 12-well plates (Corning Cat# 353043) with poly-L-lysine coated 18 mm coverslips (neuVitro Cat# GG-18-1.5-PLL) or glass-bottom 12-well plates (MatTek Corp Cat# P12G-1.5-10-F). The next day, the cells were briefly washed with ice-cold 1X PBS (R&D Systems Cat# 4870-500). Cells were fixed in 4% formaldehyde (Thermo Fisher Scientific Cat# J60401) for 15 min at room temperature, then washed three times with 1X PBS at room temperature for 5 min on an orbital shaker at 35 revolutions per minute (RPM). Cells were permeabilized with 0.1% Triton X-100 (Fisher Chemicals Cat# BP151-500) in 1X PBS for 10 min at room temperature, then washed three times with 1X PBS for 5 min on an orbital shaker. Wells were blocked with Rockland Blocking Buffer (Rockland Immunochemicals Cat# MB070) for 30 min and probed as indicated with the following primary antibodies in 1% BSA in 1X PBS-0.1% Tween-20 and its corresponding dilutions: 1:50 Alexa-Fluor 594-conjugated Vimentin (Cell Signaling Technology Cat# 7675, RRID:AB_ 2797632), Alexa-Fluor 647-conjugated ATP1A1 (Thermo Fisher Scientific Cat# MA3-928-A647, RRID:AB_2633350), and 1:100 Alexa Fluor 488-conjugated EGFR (Cell Signaling Technology Cat# 5616, RRID:AB_10691853). Coverslips were incubated in antibody dilution in a humidified chamber at 4°C overnight at minimum oscillation speed in a light-blocking container. Antibody mixtures were aspirated, then cells were washed three times with 1X PBS for 5 min on an orbital shaker in a light-blocking container. Coverslips were counterstained by adding mounting media with 4',6-diamidino-2-phenylindole (DAPI) (Vectashield Cat# H-1000-10), then placed over the DAPI drop on the microscope slide and pressed gently to remove bubbles, then dabbed to remove excess media if necessary, before sealing with nail polish. Imaging was performed using the Nikon A1 Confocal microscope with 60X objective oil immersion. Image processing was performed using the NIS Elements software.

**Flow Cytometry**

After trypsinization and neutralization with normal media, cells were filtered through 35 µm cell strainer caps (Corning Cat# 352235). A subset of cells was diluted 1:1 in Trypan blue (Invitrogen Cat# T10282) and live cells were quantified using Countess Cell Counting Slides (Invitrogen Cat# C10228) on a Countess 3 Automated cell counter (Invitrogen Cat# AMQAX2000). Approximately 500,000 cells were fixed and permeabilized by 4% formaldehyde (Thermo Fisher Scientific Cat# J60401-AP) and 90% ice-cold methanol (Sigma-Aldrich Cat# 34860), respectively. Cells were then treated with 1:50 Alexa-Fluor 488-linked EGFR (Cell Signaling Technology Cat# 5616, RRID:AB_10691853) in flow buffer for 1 hour at room temperature in a light-blocking container. Unbound antibodies were washed out and cells were resuspended in fresh flow buffer. Cells were analyzed using a BD FACSCalibur. Live cells were gated with side scatter and forward scatter. Alexa-Fluor 488-conjugated EGFR-positive cells were gated using parameters in which unstained controls had <0.1% positive cells. The Alexa-Fluor 488-linked EGFR antibody (Cell Signaling Technology Cat# 5616, RRID:AB_10691853) used for flow cytometry was selected because it binds the intracellular domain, minimizing the chances that N361A impacts binding.

***In situ* Proximity Ligation Assay (PLA) Coupled with Immunofluorescence Assay**

For each cell line, live cells were counted using a Countess 3 Automated cell counter and seeded at 200,000 cells per well in glass-bottom 12-well plates (MatTek Corp Cat# P12G-1.5-10-F) in complete media. The following day, MCF10A cells were subsequently washed with ice-cold 1X PBS (R&D Systems Cat# 4870-500) three times. 293T cells were incubated with an additional 20 ng/ml EGF for 15 minutes, then subsequently washed with ice-cold 1X PBS three times. Cells were fixed in 4% formaldehyde (Thermo Fisher Scientific Cat# J60401) for 15 min at room temperature, then permeabilized with 0.5% Triton X-100 (Fisher Chemicals Cat# BP151-500) in 1X PBS for 10 min at room temperature. Wells were then washed three times with 1X PBS at room temperature for 5 min per wash on an orbital shaker at 35 RPM, then followed by a brief wash in $ddH_2O$.

The DUOLINK *in situ* Far-Red kit (Sigma-Aldrich Cat# DUO92013) was used for the PLA. Wells were blocked with DUOLINK blocking solution with gentle oscillation for 30 min at 37°C and probed as indicated with the following primary antibodies in DUOLINK Antibody Diluent and its corresponding dilutions: 1:100 anti-rabbit monoclonal Her2/ErbB2 (Cell Signaling Technology Cat# 2165, RRID:AB_10692490) and 1:100 anti-mouse monoclonal EGFR (Genetex Cat# GTX628887, RRID:AB_2888064). To avoid possible interactions with the extracellular N361 region, these antibodies each bind the intracellular domain of their target: residues surrounding

Tyr1248 of Her2 and the C-terminal region of human EGFR. Well plates were incubated in primary antibody dilution as mentioned above in a humidified chamber at 4°C overnight at minimum oscillation speed in a light-blocking container. Three washes using DUOLINK *in situ* wash buffers for fluorescence, Buffers A and B (Sigma-Aldrich Cat# DUO82049), were performed at room temperature on an orbital shaker at minimum oscillation speed for 5 min each. Wells were probed with DUOLINK PLA probe solution, using equal amounts of PLA probe 1:5 Donkey anti-Mouse MINUS (Sigma-Aldrich Cat# DUO92004, RRID:AB_2713942) and 1:5 Donkey anti-Rabbit PLUS (Sigma-Aldrich Cat# DUO92002, RRID:AB_2810940) diluted in DUOLINK Antibody diluent for an hour in a humidified chamber at 37°C. Wells were then washed twice with Buffer A for 5 min. Ligation was then performed by incubating wells with DUOLINK ligation solution (1:40 1 U/ul ligase in 1X ligation buffer in high-purity $H_2O$), followed by DUOLINK amplification solution (1:80 10 U/ul polymerase in 1X amplification FarRed buffer in high-purity $H_2O$). Ligation of MINUS and PLUS probes for each well was performed by incubation with DUOLINK *in situ* ligation solution for 30 min in a humidified chamber at 37°C, then washed twice with Buffer A for 2 min. Amplification of signal in each well was performed by incubation with DUOLINK *in situ* amplification solution for 100 min in a humidified chamber at 37°C, then washed initially with Buffer B for 10 min and then washed again with a 1:100 dilution of the same buffer for 1 min. All steps after amplification were performed in a light-blocking container covering the well plate. All incubation steps at 37°C were performed without any shaking or oscillation, and incubation steps at 4°C were performed on an orbital shaker at the aforementioned speed. Each well was then incubated with two drops per mL of ActinRed 555 ReadyProbe fluorescence dye (Sigma-Aldrich Cat# R37112), then washed twice with 1X PBS at room temperature on an orbital shaker at the aforementioned speed (35 RPM). Finally, wells were counterstained with DUOLINK *in situ* mounting media with DAPI (Sigma-Aldrich Cat# DUO82040) and incubated at room temperature for 15 min.

Imaging was performed using the Nikon A1 Confocal microscope with 60X objective oil immersion. Image processing was performed using the included NIS Elements software. Cell boundaries were defined using ActinRed channel. Quantification of mean pixel intensity per cell line was determined by CellProfiler Image Analysis Software (RRID:SCR_007358).[40] The same settings were applied to each of the 5 replicates per condition.

#### Figure Preparation

Figures were prepared using Adobe Illustrator Creative Cloud 2023.

### Supplemental Figure 1

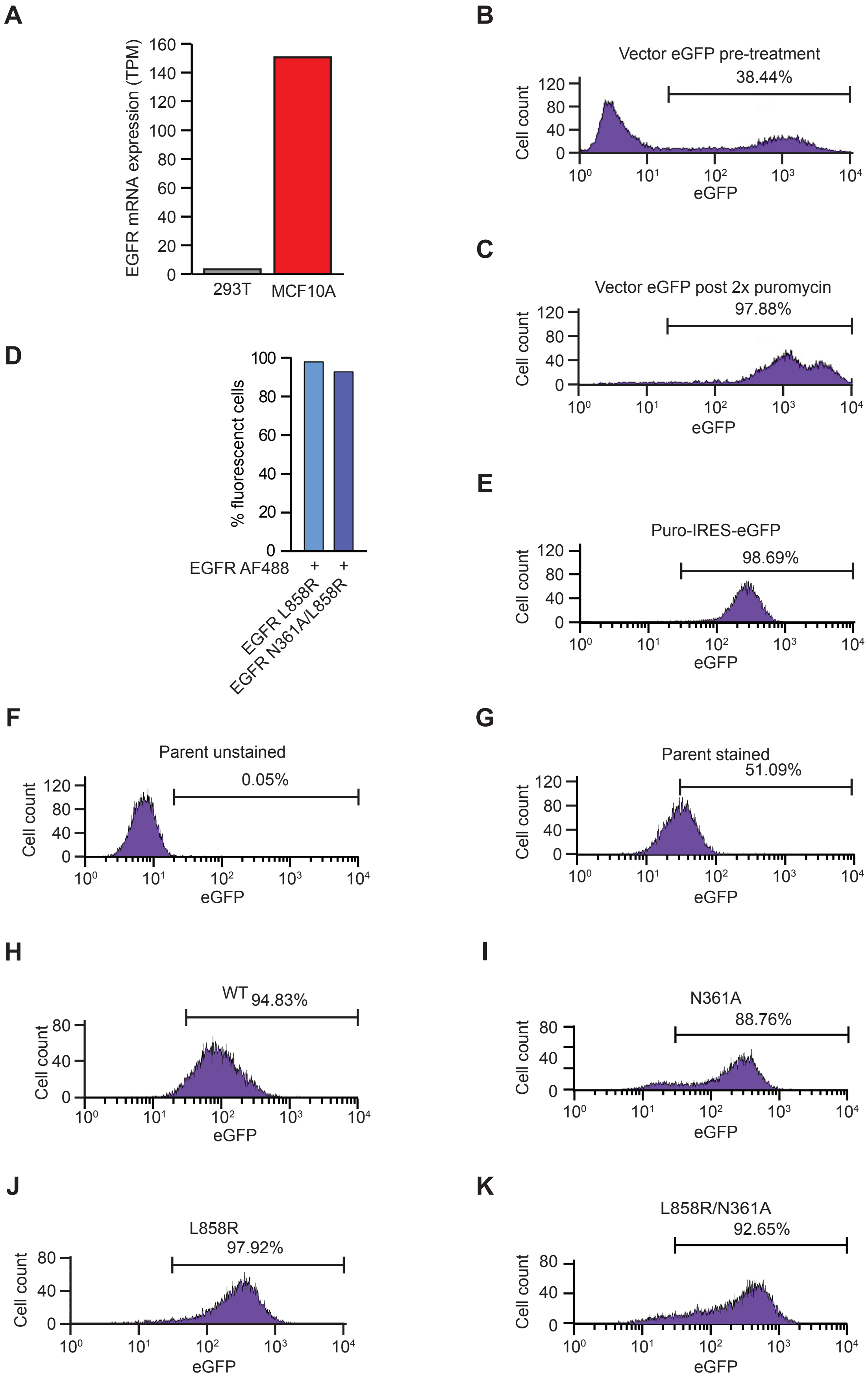

**Supplementary Figure 1. EGFR overexpression constructs in cells. A,** mRNA expression of EGFR in HEK-293T and MCF10A cells from single-cell RNAseq data. Adapted from Human Protein Atlas.[67] **B-C,** Histogram of flow cytometry data showing eGFP in positive control MCF10A cells overexpressing fluorescent puro-IRES-eGFP before and after two rounds of puromycin selection. **D,** Percentage of fluorescent MCF10A cells, overexpressing either EGFR L858R or a single construct containing two mutations EGFR N361A/L858R, from flow cytometry labeled with anti-EGFR antibody conjugated to Alexa-Fluor 488, or empty fluorescent vector puro-IRES-GFP. **E-K**, Histograms of flow cytometry data of cells stained with a fluorescent EGFR-AF488 antibody or control unstained cells. Shown are **(E)** unstained parental MCF10A cells expressing puro-IRES-eGFP, **(F)** unstained parental cells, **(G)** stained parental cells, **(H)** EGFR WT cDNA, **(I)** EGFR N361A cDNA, **(J)** EGFR L858R cDNA, or **(K)** a single cDNA construct containing two mutations EGFR N361A/L858R.

### Supplemental Figure 2

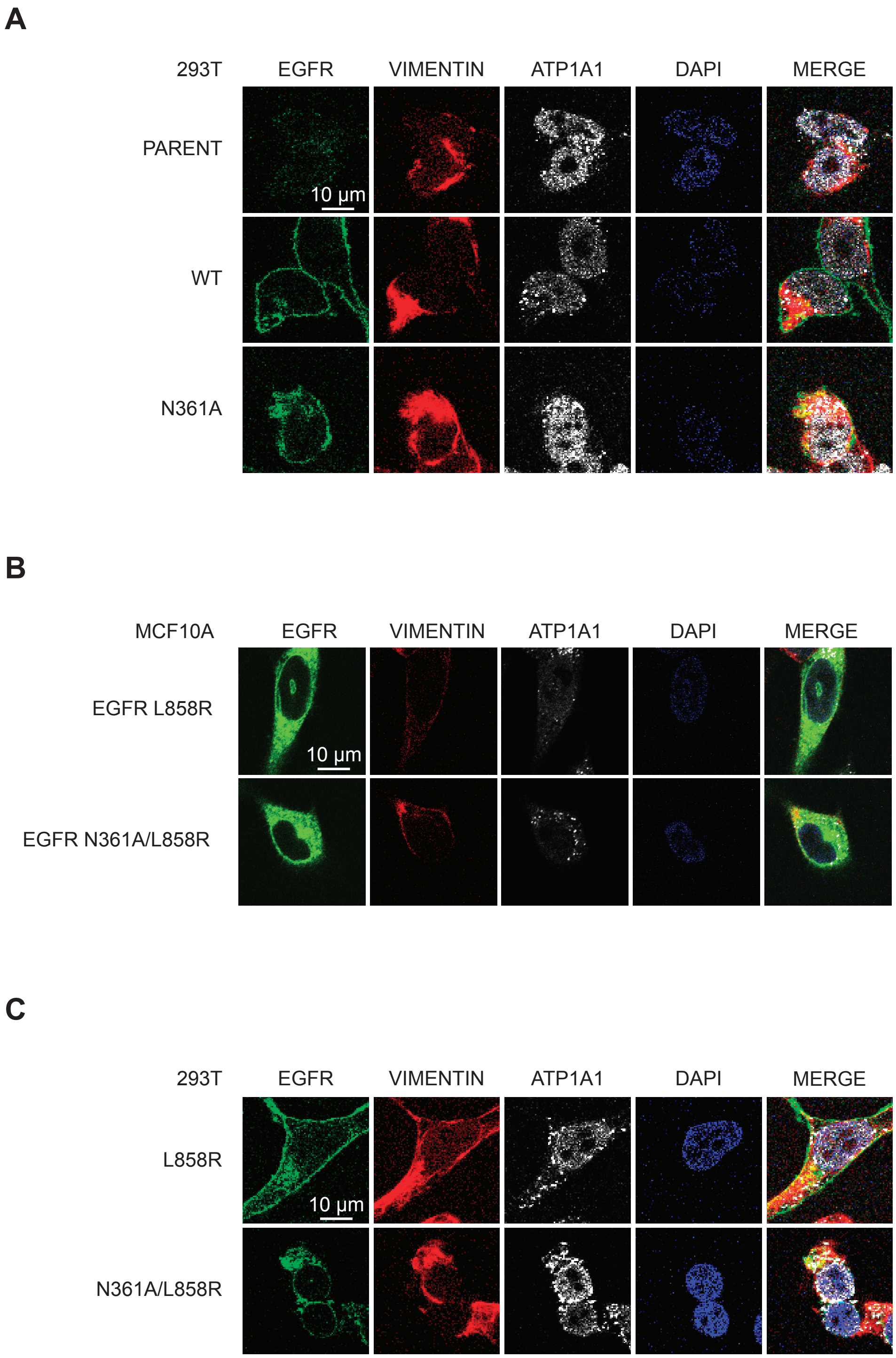

**Supplementary Figure 2. A,** Representative immunofluorescent microscopy images of parental 293T cells or 293T cells overexpressing cDNAs of EGFR wild-type (WT) or EGFR N361A. **B,** Representative immunofluorescent microscopy images of 293T cells overexpressing cDNAs of EGFR L858R or a single construct containing two mutations EGFR N361A/L858R. **C,** Representative immunofluorescent microscopy images of MCF10A cells overexpressing cDNAs of EGFR L858R or a single construct containing two mutations EGFR N361A/L858R.

### Supplemental Figure 3

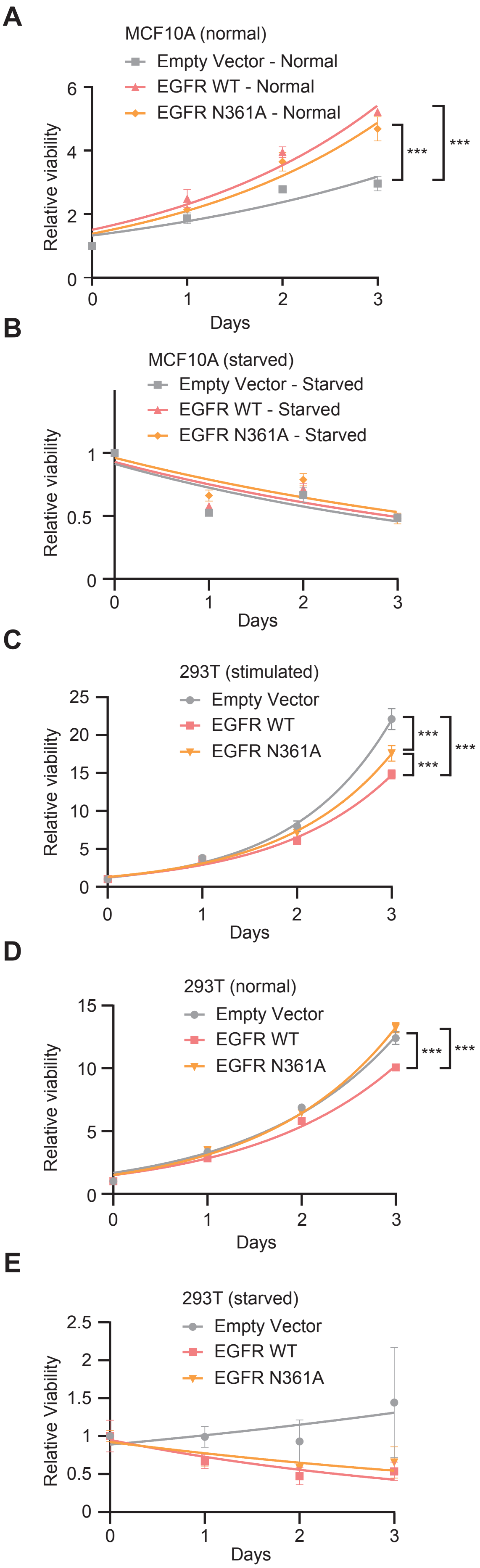

**Supplementary Figure 3. A-B,** Time courses of CTG relative viability of parental MCF10A cells or MCF10A cells overexpressing cDNAs of EGFR WT or EGFR N361A in cells in **(A)** normal media for **(B)** starved media, **C-E,** Time courses of relative viability of parental 293T cells or 293T cells overexpressing cDNAs of EGFR WT or EGFR N361A in **(C)** stimulated media, **(D)** normal media, **(E)** starved media. ns = not significant, * $p < 0.05$; *** $p < 0.001$, $n = 3$.

Supplemental Figure 4

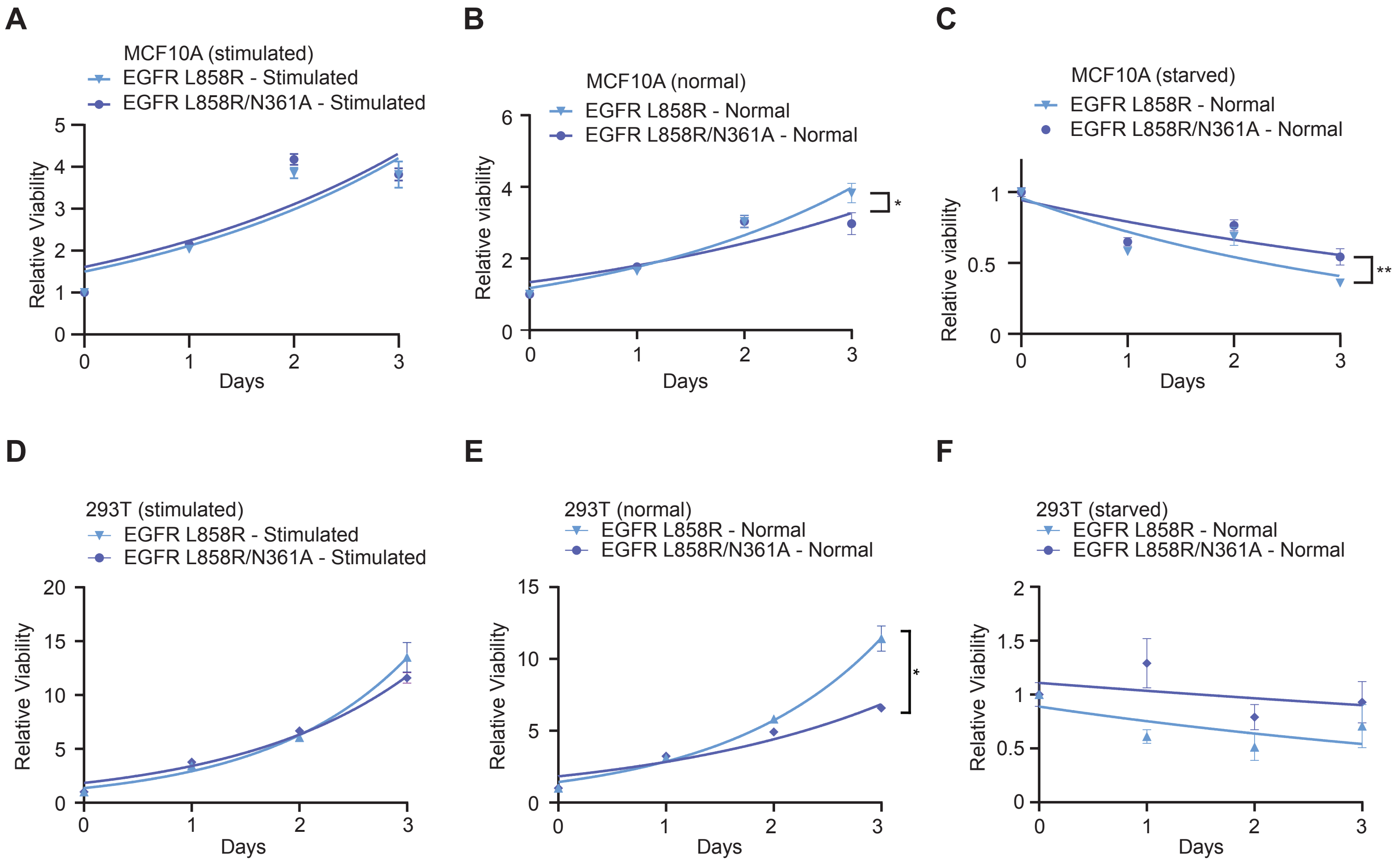

**Supplementary Figure 4. A-C,** Time courses of relative viability of MCF10A cells overexpressing cDNAs of EGFR L858R or a single construct containing two mutations EGFR N361A/L858R in **(A)** stimulated media, **(B)** normal media, **(C)** starved media. **D-F,** Time courses of relative viability of 293T cells overexpressing cDNAs of EGFR L858R or a single construct containing two mutations EGFR N361A/L858R in **(D)** stimulated media, **(E)** normal media, **(F)** starved media. ns = not significant, * $p < 0.05$; *** $p < 0.001$, n = 3.

Supplemental Figure 5

A

B

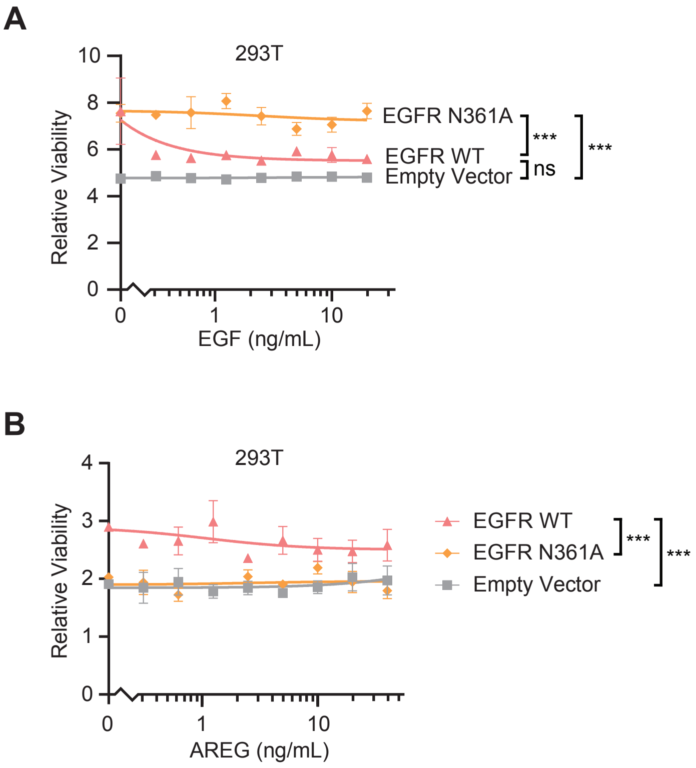

**Supplementary Figure 5. A-B,** Dose course of relative viability of parental 293T cells or 293T cells overexpressing cDNAs of EGFR WT or EGFR N361A upon stimulation by **(A)** EGF or **(B)** AREG for 72 hours, both measured by CellTiter-Glo. ns = not significant, * $p < 0.05$; *** $p < 0.001$, $n = 3$.

### Supplemental Figure 6

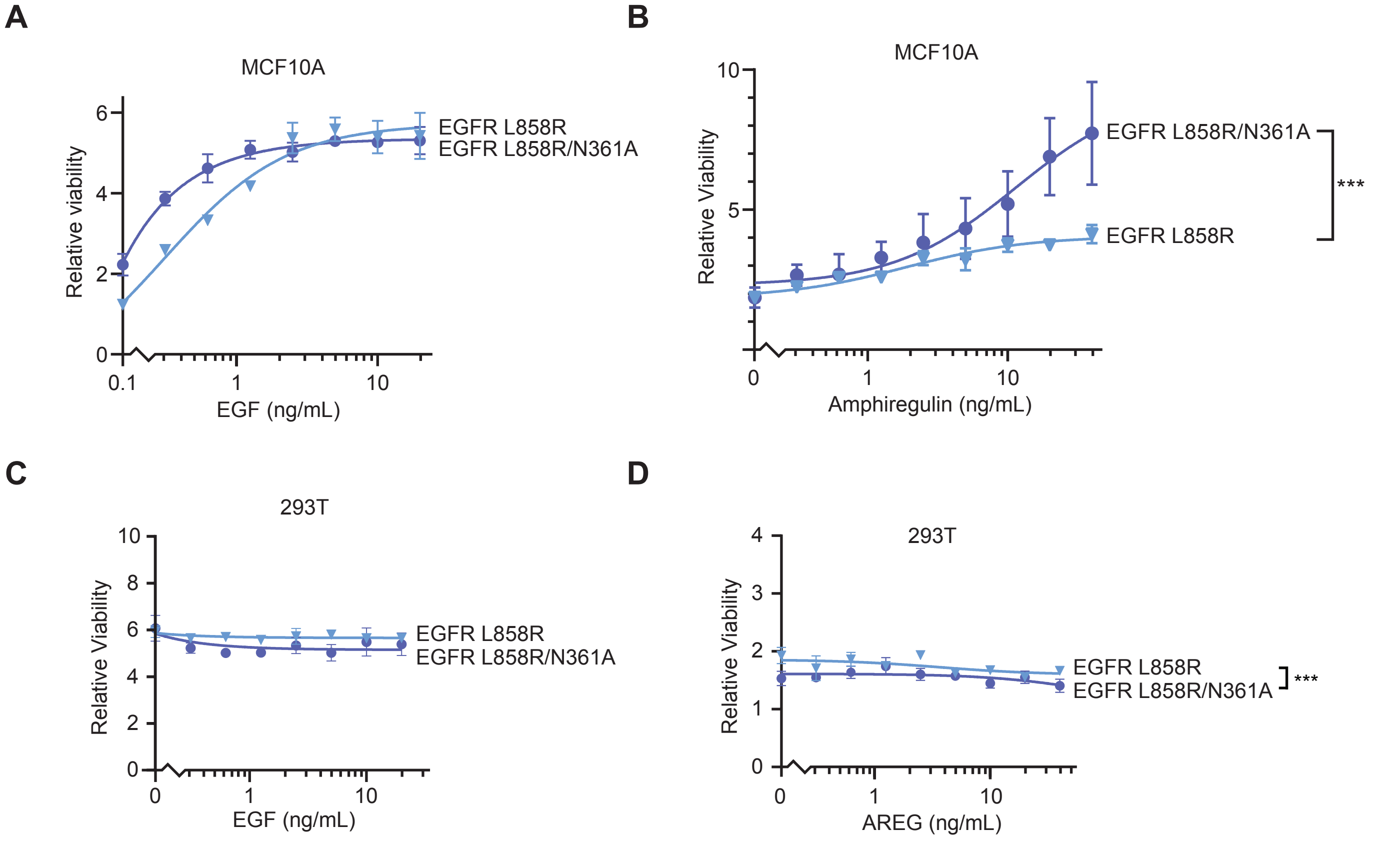

**Supplementary Figure 6. A-B,** Dose course of relative viability of MCF10A cells overexpressing cDNAs of EGFR L858R or a single construct containing the two mutations EGFR N361A/L858R upon stimulation by **(A)** EGF or **(B)** AREG for 72 hours, both measured by CTG. **C-D,** Dose course of relative viability of 293T cells overexpressing cDNAs of EGFR L858R or a single construct containing two mutations EGFR N361A/L858R upon stimulation by **(C)** EGF or **(D)** AREG for 72 hours, both measured by CTG. ns = not significant, * $p < 0.05$; *** $p < 0.001$, $n = 3$.

### Supplemental Figure 7

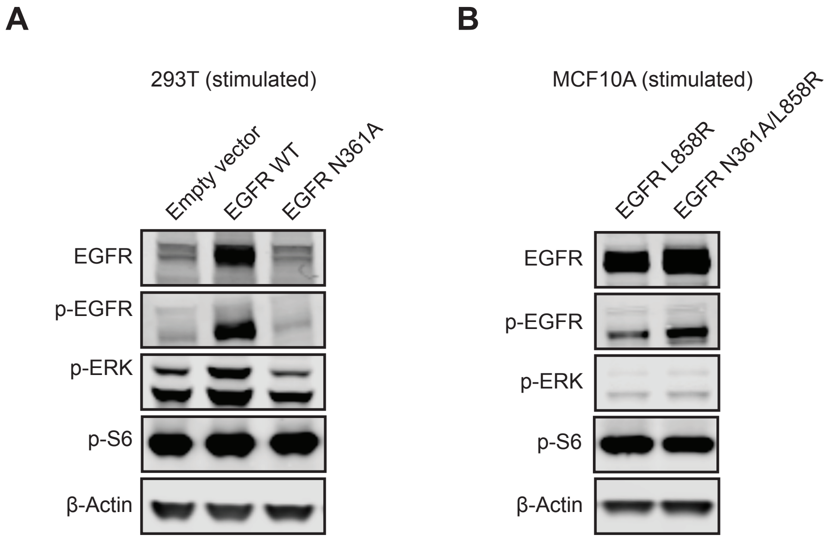

**Supplementary Figure 7. A,** Immunoblots of whole cell lysates of MCF10A cells overexpressing cDNAs of EGFR L858R, or a single construct containing two mutations EGFR N361A/L858R stimulated with an additional 20 ng/mL of EGF for 15 minutes. **B,** Immunoblots of whole cell lysates of parental 293T cells or 293T cells overexpressing cDNAs of EGFR WT, EGFR N361A stimulated with an additional 20 ng/mL of EGF for 15 minutes. **C,** Immunoblots of whole cell lysates of 293T cells overexpressing cDNAs of EGFR L858R, or a single construct containing two mutations EGFR N361A/L858R stimulated with an additional 20 ng/mL of EGF for 15 minutes.

### Supplemental Figure 8

A

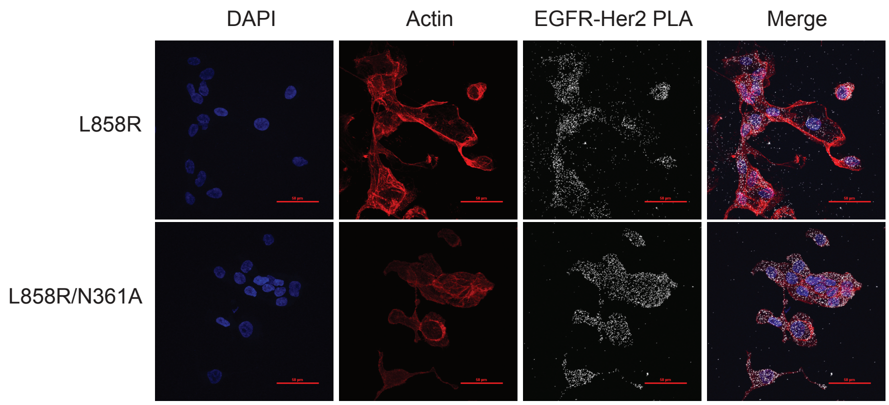


B

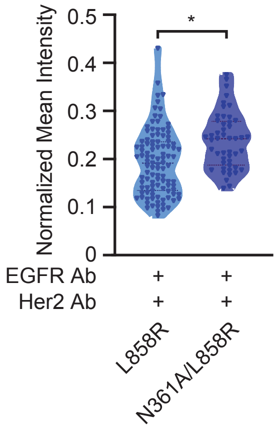


C

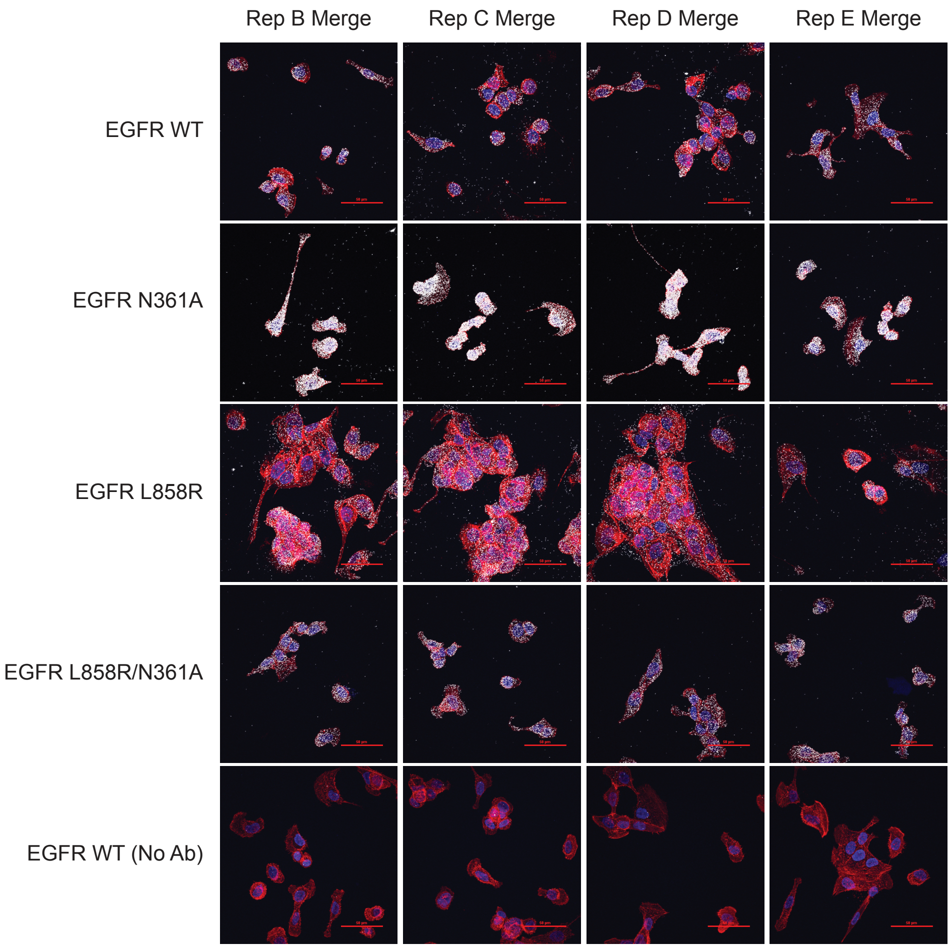

**Supplementary Figure 8. A,** Immunofluorescent images from *in situ* proximity ligation assays (PLA) measuring colocalization of HER2 and EGFR in MCF10A cells overexpressing cDNAs of EGFR L858R, or a single construct containing two mutations EGFR N361A/L858R. **B,** Representative quantification of mean pixel intensity of cellular regions defined by rhodamine Actin stain. * $p < 0.05$, $n = 5$. **C,** Images of additional PLA replicates of MCF10A cells overexpressing the indicated cDNAs or negative control with no antibodies. Including Figure 3A, $n = 5$.

### Supplemental Figure 9

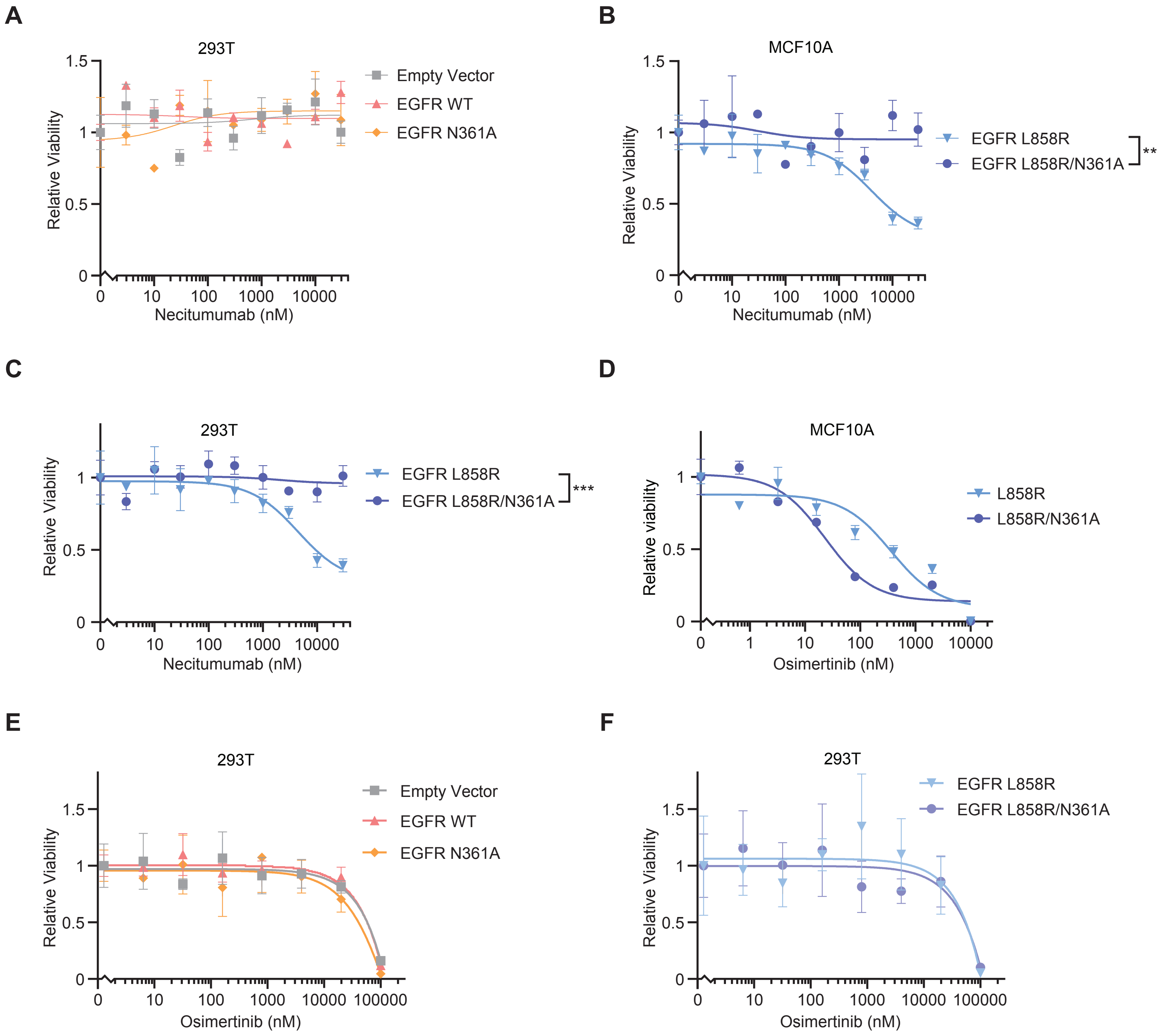

**Supplementary Figure 9.** Dose courses of relative viability of the indicated cells overexpressing the indicated cDNAs upon treatment with the indicated drug for 72 hours, measured by CellTiter-Glo. **A,** 293T cells overexpressing EGFR WT, EGFR N361A, or empty vector treated with necitumumab. **B,** MCF10A cells overexpressing EGFR L858R or EGFR N361A/L858R treated with necitumumab. **C,** 293T cells overexpressing EGFR L858R or EGFR N361A/L858R treated with necitumumab. **D,** MCF10A cells overexpressing EGFR L858R or EGFR N361A/L858R treated with osimertinib. **E,** 293T cells overexpressing EGFR WT, EGFR N361A, or empty vector treated with osimertinib. **F,** 293T cells overexpressing EGFR L858R or EGFR N361A/L858R treated with osimertinib. *** $p < 0.001$, $n = 3$.
